# Pitfalls and windfalls of detecting demographic declines using population genetics in long-lived species

**DOI:** 10.1101/2024.03.27.586886

**Authors:** Meaghan I. Clark, Sarah W. Fitzpatrick, Gideon S. Bradburd

## Abstract

Detecting recent demographic changes is a crucial component of species conservation and management, as many natural populations face declines due to anthropogenic habitat alteration and climate change. Genetic methods allow researchers to detect changes in effective population size (N_e_) from sampling at a single timepoint. However, in species with long lifespans, there is a lag between the start of a decline in a population and the resulting decrease in genetic diversity. This lag slows the rate at which diversity is lost, and therefore makes it difficult to detect recent declines using genetic data. However, the genomes of old individuals can provide a window into the past, and can be compared to those of younger individuals, a contrast that may help reveal recent demographic declines. To test whether comparing the genomes of young and old individuals can help infer recent demographic bottlenecks, we use forward-time, individual-based simulations with varying mean individual lifespans and extents of generational overlap. We find that age information can be used to aid in the detection of demographic declines when the decline has been severe. When average lifespan is long, comparing young and old individuals from a single timepoint has greater power to detect a recent (within the last 50 years) bottleneck event than comparing individuals sampled at different points in time. Our results demonstrate how longevity and generational overlap can be both a hindrance and a boon to detecting recent demographic declines from population genomic data.

## 1. Introduction

Modern-day wild populations are experiencing declines and bottleneck events from an unprecedented array of threats, including climate change, habitat fragmentation, and disease (Haddad et al., 2015; McCarty, 2001; Pimm et al., 2014). Population responses to these threats are often species-specific and context-dependent (Debinski & Holt, 2000; Fitzpatrick et al., 2023); for example, populations of species with specialized habitat requirements are more likely to experience sharp demographic declines when their habitat is disturbed than populations of generalist species. Populations that have undergone dramatic declines are at risk of extirpation due to environmental and demographic stochasticity and have a higher risk of experiencing inbreeding depression compared to populations that have been historically small (D. Charlesworth & Willis, 2009; Lande, 1988). The timely detection of recent population declines is essential for effective conservation intervention and ensuring the persistence of imperiled populations.

Although critical, detecting recent changes (within ∼50 years) in population size is a long-standing challenge in conservation biology and evolutionary theory (Peery et al., 2012). Field-based methods to detect population size changes (Chapman, 1954; Gazey & Staley, 1986), such as mark-recapture models and long-term population monitoring, can be used to generate precise estimates of census population size (N_c_) and other demographic trends over time (Congdon et al., 1993). However, estimation of N_c_ from field methods is labor-intensive, difficult to conduct for cryptic species, and cannot provide estimates of historic population size.

Genetic methods to detect changes in population size are often faster and logistically easier than field-based methods, and become more affordable and informative every year (Hohenlohe et al., 2021). Rather than measure N_c_ directly, these methods estimate changes in effective population size (N_e_) over time from genetic data, often from a single sampling point. N_e_ represents the number of individuals in an idealized Wright-Fisher population with the same genetic behavior (e.g., same rate of genetic drift, or same amount of genetic diversity) as the focal empirical population (Charlesworth, 2009; Waples, 2022). Because of many factors, including changes in population size and the magnitude of heterogeneity in reproductive success, N_e_ is usually smaller than N_c_ for a given empirical population (Fisher, 1923; Wright, 1931). Current methods of detecting demographic trends from genetic data are quite accurate when inferring ancient demographic trends (> ∼100 generations in the past) but have low power to capture trends associated with recent events (within the last ∼100 generations) without large amounts of sequencing data and/or individuals sampled (Antao et al., 2011; Beichman et al., 2018; Clark et al., 2023; Reid & Pinsky, 2022). Repeated temporal sampling can be a powerful tool for detecting genetic patterns associated with a decline, especially when sample size is limited (Antao et al., 2011; Clark et al., 2023), but such sampling is unlikely to exist for most populations of concern.

Using genetic methods to detect demographic declines is particularly fraught in species with long lifespans and overlapping generations because the loss of genetic diversity associated with a decline happens on a timescale determined in part by generation time (Felsenstein, 1971; Gargiulo et al., 2024 preprint; Hill, 1972; Kuo & Janzen, 2003). Generational overlap slows the rate of genetic drift, as individuals contribute to multiple future cohorts. Long lifespan also reduces the efficacy of temporal sampling because more years will have to elapse before an appreciable change in N_e_, as measured by genetic diversity, is detected. Generational overlap and long lifespan, which are usually correlated across the tree of life (Jones et al., 2018), generate a lag between a demographic decline and the resulting decrease in N_e_ and genetic diversity (Nunney, 1993). Survivors of a decline persist in the population and can reproduce and contribute to multiple future age cohorts, slowing the loss of genetic variation due to genetic drift compared to organisms with a short lifespan and less generational overlap. This phenomenon reduces the power of demographic inference and bottleneck tests to detect recent population declines in long-lived species. In multiple empirical systems of long-lived species, including the copper redhorse (Lippé et al., 2006), ivory gull (Charbonnel et al., 2022), spotted turtles (Davy & Murphy, 2014), black abalone (Wooldridge et al., 2024), and white-tailed eagles (Hailer et al., 2006), researchers have had difficulty detecting population bottlenecks with genetic data despite strongly suspected or documented decline in N_c_. Further, simulation studies have found lower accuracy in detecting demographic declines with increased generation time (Bradke et al., 2021; Reid & Pinsky, 2022).

The longevity of a species can represent a hurdle for detecting demographic declines with genetic data, but it also presents an opportunity. First, generational overlap should decrease the rate at which variants are lost due to genetic drift after a population decline, giving conservation practitioners more time to intervene before inbreeding depression and loss of adaptive potential send a population into the extinction vortex (Gilpin & Soulé, 1986). Second, surviving individuals in the post-decline population retain genetic diversity representative of the pre-decline gene pool, whereas younger individuals born into the post-decline population come from a smaller set of possible parents and may have lower genetic diversity (Figure 1A), indicating that a relationship between individual age and genetic diversity might be an early indicator of a population decline.

**Figure 1.**
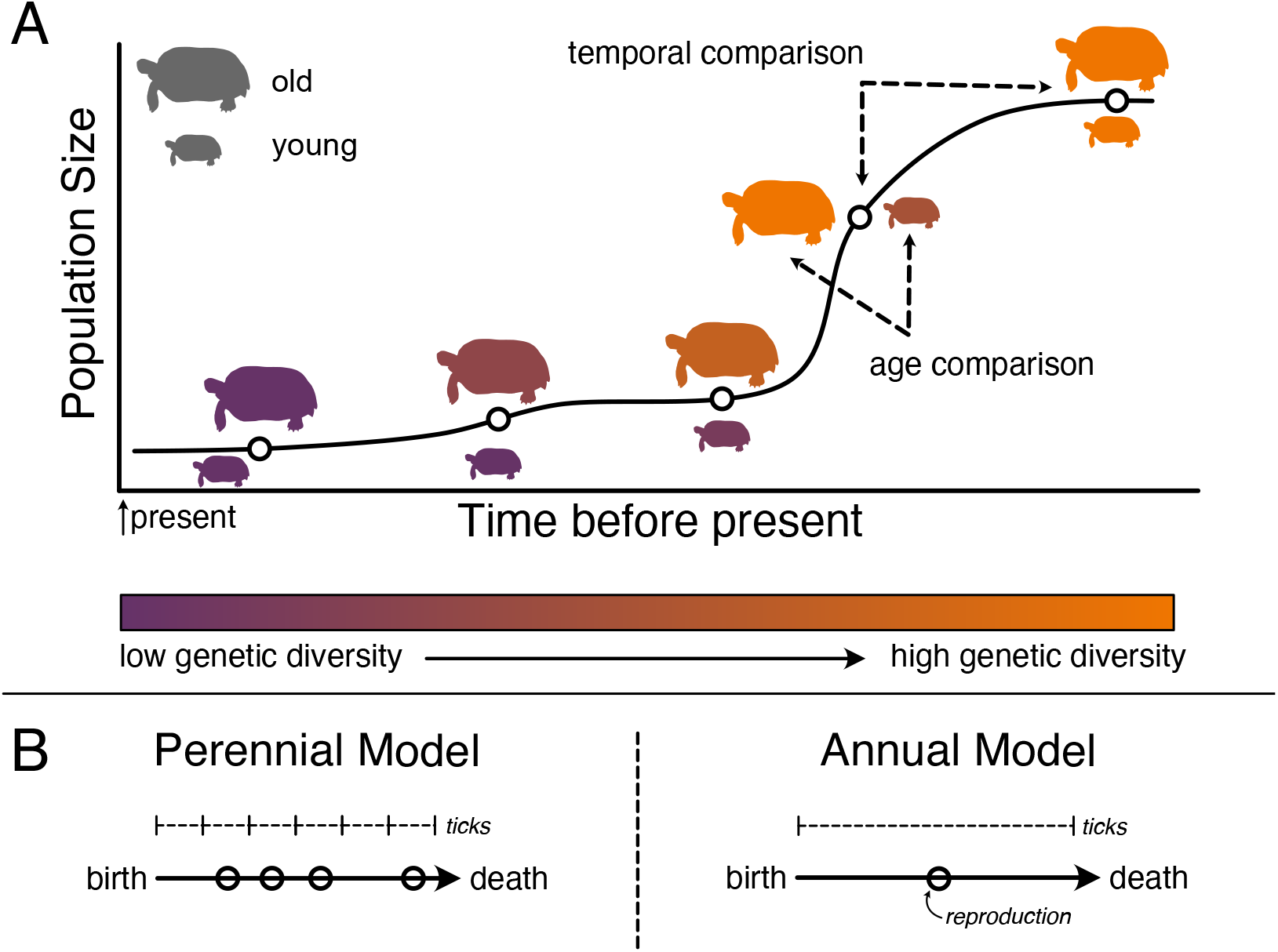
(A) In species with overlapping generations, older individuals born before the bottleneck will have genomes representative of a larger population size. Figure shows a population of tortoises that have undergone a population decline. At sampling points (open circles) before the decline, old and young turtles have similar levels of genetic diversity. During the decline, old individuals who were born before the decline have high levels of diversity compared to younger individuals, born after the decline from a limited set of possible parents. At some point after the decline, the population reaches a new equilibrium level of diversity, and old and young individuals have similar levels of genetic diversity. Dotted lines represent comparisons presented in this study: comparisons between adjacent timepoints (temporal) and comparisons between different age bins (age). (B) Schematics depicting life cycles in the annual and perennial models. In the annual model, each individual lives for one tick of the simulation and reproduction occurs once. In the perennial model, each individual lives for multiple ticks, with a probability of mortality (*p*) in each tick and has opportunities to reproduce in each tick.

Several empirical studies have compared genetic diversity between older and younger individuals in long-lived species to look for evidence of population decline or the genetic consequences of known decline and fragmentation, with mixed results (Kettle et al., 2007; Labonne et al., 2016; Pereira et al., 2023; Schmidt et al., 2018; Vranckx et al., 2012; Yineger et al., 2014). A meta-analysis by Vranckx et al. (2012) found that studies comparing genetic diversity between adult and sapling trees in fragmented habitat overall found higher allelic richness in adult trees. On the other hand, Schmidt et al. (2018) did not find a relationship between heterozygosity and age in the long-lived Australian lungfish (*Neoceratodus forsteri*), despite habitat degradation and suspected decline. The development of new methods for estimating individual age non-invasively, for example using telomere length (Haussmann & Vleck, 2002), DNA methylation levels (Nakamura et al., 2023), and radiocarbon dating (Fallon et al., 2019), expands the applicability of age comparisons in long-lived organisms of high conservation concern. However, to understand the utility of using genetic differences and age structure within contemporary populations as evidence of demographic decline, we need to investigate the relationship between age and genetic patterns in a system where the true timing and severity of the decline is known.

In this study, we use simulations to ask under what demographic and life history conditions it is useful to treat the genomes of older individuals as pseudo-temporal sampling compared with those of younger individuals to learn about recent changes in N_e_. To fully understand the genetic dynamics involved in population declines in long-lived species, we track the fate of simulated individuals in populations implemented in SLiM (Haller & Messer, 2019). We simulate organisms with varying longevity, characterize the genetic patterns associated with recent demographic changes and test if sampling from different age groups can successfully identify recent bottlenecks of varying strengths. We compare these results to temporal sampling that does not consider age information and to simulations without overlapping generations to evaluate how longevity can be both a hindrance and a boon in detecting recent demographic declines from population genomic data. Our simulations reveal that comparisons between age groups do have power to detect severe bottlenecks in species with long lifespans, highlighting how, and under what conditions, age information can be leveraged to learn about recent changes in N_e_.

## 2. Materials and Methods

### 2.1. Simulation set-up

Demographic scenarios were modeled using simulations tracking the fate of individuals in a population in SLiM v.3.7.1 (Haller & Messer, 2019). We built two non-Wright-Fisher simulations, one representing a species with a long lifespan (average age in population ≥ 2 simulation ticks) and overlapping generations (Figure 1B, “perennial simulation”), and one representing a species with a short lifespan with non-overlapping generations (Figure 1B, “annual simulation”). Both simulations contained instructions for reproduction and mortality events, which occur at each time-step, or tick, of the simulation. Ticks can be thought of as years. Individuals in both models were hermaphroditic and selfing was prohibited. Each simulation is detailed below.

#### 2.1.1 Perennial simulation

The perennial simulation mimics the life history of species such as trees and turtles––organisms that reproduce on an annual cycle and have considerable generational overlap with long lifespans. In the perennial simulation, individuals survive for multiple reproductive ticks, with *p* representing the probability of mortality in each tick. The average lifespan in the model is 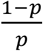. The lifespan of individuals in the model can be manipulated in a predictable manner by changing the value of *p*. Individuals have a constant mortality probability across all age classes.

The census population size (N_c_) is an emergent property of the simulation and can be modified using the parameters 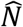 (desired equilibrium population size) and *K* (the “carrying capacity” of the model). Carrying capacity *K* is defined as 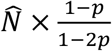. A static value of 1 – *p* determines if an individual lives or dies within a tick of the simulation. In each tick of the model, offspring are produced from individuals who are one tick old and older. The probability of a focal individual having offspring is defined as either zero or 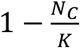, whichever is larger. If N_c_ is less than *K*, the probability of reproduction will be greater than zero and if N_c_ is more than *K*, the probability of reproduction is zero. Thus, population size is maintained at approximately 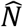 through reproduction. A second parent individual is drawn at random from the pool of individuals one tick old and older, and a single offspring is generated per pair. The perennial model implements an instantaneous bottleneck event by changing *K* to *B*, where 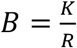 and *R* is an integer representing bottleneck severity, when calculating the probability of offspring, and by scaling fitness by 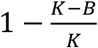 in the bottleneck tick.

#### 2.1.2 Annual simulation

The annual simulation mimics the life history of organisms such as univoltine mosquitoes, butterflies, and annual plants––organisms that have a single reproductive opportunity before dying. In the annual simulation, individuals survive for a single reproductive cycle. Census population size (N_c_) is still an emergent property of the simulation, modified using 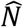 and *K*. In this simulation, *K* is equal to 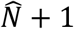 to maintain N_c_ at 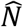.

In each tick of the simulation, offspring are produced from the previous generation of individuals (all one tick of age). The probability of an individual producing offspring is defined as 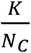 (in practice, this ratio was 1 at almost every cycle). The number of offspring produced per focal individual is drawn from a Poisson distribution with a lambda equal to the probability of producing offspring. The model implements a bottleneck event by changing *K* to *B* where 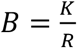 (where *R* again represents bottleneck severity) when calculating the probability of offspring.

### 2.2. Simulation implementation and analysis

All perennial simulations ran for a burn-in period of approx. 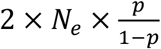 ticks (Table 1) to ensure equilibrium age distributions. The annual simulations were run with a burn-in of 100,000 ticks. After the burn-in period, a bottleneck event occurred, where N_c_ was reduced by a factor *R*. All simulations were run for 500 ticks after the bottleneck event. SLiM recorded genealogical information for 500 randomly selected individuals every 50 ticks for 200 ticks before the bottleneck, every 5 ticks for 50 ticks after the bottleneck, and every 50 ticks for an additional 450 ticks until the end of the simulation for a total of 24 sampling timepoints. We recorded age (in ticks) for every sampled individual. At the end of the simulations, we output the genealogical history of sampled individuals as tree sequences, a computationally efficient file format for storing information about genealogical networks and genetic variation within a sample (Kelleher et al., 2018; Lewanski et al., 2024).

**Table 1.**
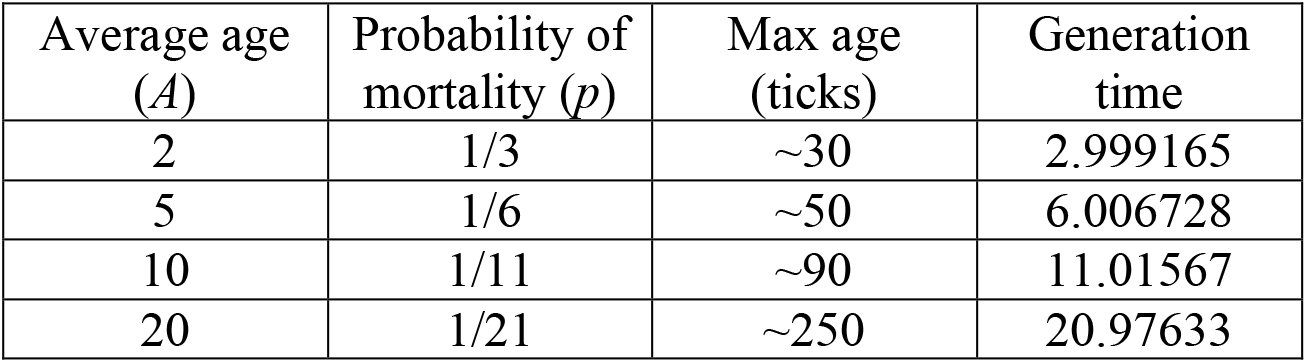
Average age, probability of mortality, maximum age in ticks, and generation time of perennial simulations.

| Average age<br>($A$) | Probability of<br>mortality ($p$) | Max age<br>(ticks) | Generation<br>time |
| --- | --- | --- | --- |
| 2 | 1/3 | ~30 | 2.999165 |
| 5 | 1/6 | ~50 | 6.006728 |
| 10 | 1/11 | ~90 | 11.01567 |
| 20 | 1/21 | ~250 | 20.97633 |

We ran 100 replicates of the annual simulation for each of three values of *R* (2, 10 or 100). We ran 100 replicates of the perennial simulation with four different values of *p*, and three different values of *R* (Table 1, 1200 simulations total). Overlapping generations decreases N_e_ more than expected by Wright-Fisher expectations because fewer individuals are born into the population each reproductive cycle (assuming constant population size), and there is higher variance in lifetime reproductive success (Hill, 1972). To achieve similar pre-bottleneck N_e_ between simulations, we raised N_c_ in the perennial models such that N_e_ of each model was equivalent (Hill, 1979).

### 2.3. Tree sequence processing

To efficiently analyze all simulations, we used tree sequences from SLiM to generate genetic data for downstream analyses. For tree sequences with multiple roots in which coalescence had not yet occurred, we simulated coalescence backwards-in-time (recapitation) using msprime v.1.0.2 (Baumdicker et al., 2022; Kelleher et al., 2016) and pyslim v.1.0.4 (Haller et al., 2019). To account for generation times greater than 1 during recapitation, we scaled N_e_ and recombination rate by generation time. Recapitation also ensured that simulations reached equilibrium levels of genetic diversity. We then overlaid neutral mutations on the tree sequence using pyslim with a mutation rate of 1e^-8^. Although generation time is correlated with mutation rate in empirical systems (Bergeron et al., 2023), we chose a fixed mutation rate for all simulations to better assess the impact of lifespan alone on genetic patterns. For perennial models, the mutation rate was divided by the average generation time (as in Battey et al., 2020). To calculate generation time, we ran simplified versions of the perennial model for ten replicates per *p* value, with no mutations or bottleneck events, and recorded the average age of parents. Generation time was approximately 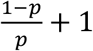 (Table 1), reflecting that individuals did not reproduce until one cycle of age.

### 2.4. Detecting declines

There are a variety of bottleneck detection methods available for reduced-representation and whole genome-sequencing data. In this paper, we focus on exploring the use of age information in improving detection, rather than testing commonly used demographic modeling approaches, which enabled us to test a wider array of parameters within computational constraints. We compare age sampling from within a single timepoint to temporal sampling using two commonly used summary statistics: Wu and Watterson’s theta, (*θ*_*W*_) and pi (*π*). *θ*_*W*_ is based on the number of segregating sites in the sample, a metric that is sensitive to changes in population size and is generally comparable across groups with different sampling sizes (Peery et al., 2012; Watterson, 1975). *π* or nucleotide diversity, is based on pairwise nucleotide differences between a set of samples (Nei & Li, 1979). Because segregating sites are lost due to genetic drift during a bottleneck event, we expect *θ*_*W*_ to be more reactive to simulated bottlenecks compared to *π*. Both *θ*_*W*_ and *π* represent the amount of genetic diversity in a population, and directly reflect long-term N_e_ (Waples, 2022).

Using tree sequences for all parameter combinations and replicates, we calculated *θ*_*W*_ and *π* at each sampling timepoint using four sampling schemes: (1) all individuals sampled to track trends in genetic diversity over time, and (2) subsamples (“bins”) containing the oldest and youngest individuals in the sample. We defined bins based on individuals whose age at a given timepoint fell above the 90^th^ quantile (“old bin”), and then selected the same number of the youngest individuals (“young bin”) for a given combination of simulation parameters and timepoints. (3) We repeat binned sampling such that that the “old bin” contained randomly sampled individuals whose age fell above the 50^th^ quantile, keeping sample size consistent with the previous sampling scheme. The binned sampling schemes represents a scenario where exact age information is not available, but individuals are able to be roughly categorized based on age, with varying degrees of certainty. Finally, (4) individuals selected at random from the sample, regardless of age, with the same sample size as binned sampling, to be used for temporal tests.

With imperfect age information, we asked if the difference in a genetic summary statistic between age bins is significant. To assess significance, we used a non-parametric permutation approach. At each timepoint, sampled individual ages were randomly shuffled, and individuals were then binned according to the binned sampling schemes described above. We calculated *π* and *θ*_*W*_ for each permuted bin for 100 permutations and calculated the difference between the two groups for each permutation to create a distribution of differences. We then calculated the one-tailed 95% quantile of that distribution and recorded if the observed difference between bins (sampling schemes 2 and 3) lay outside that boundary. To represent temporal sampling, we compared summary statistics between randomly sampled individuals at a timepoint to the previous timepoint (either 50 or 5 ticks in the past). We did 100 permutations where individuals were randomly sampled from the present and past timepoints, calculated the difference in *θ*_*W*_ and *π* between permutations and assessed if the observed difference (sampling scheme 4) between timepoints lay beyond the 95% quantile of the permuted distribution.

## 3. Results

### 3.1. The impact of lifespan and overlapping generations on genetic patterns

The rate at which genetic diversity metrics (*θ*_*W*_ and *π*) were lost after the bottleneck event decreased with increasing lifespan and corresponding generational overlap (Figure 3). Fifty ticks after the bottleneck decreased population size by a factor of 10, *θ*_*W*_ of all sampled individuals had decreased by 21.6% when average lifespan was 1 (annual simulation), 11.7% when average lifespan was 2, 7.2% when average lifespan was 5, 4.4% when average lifespan was 10, and 2.5% when average lifespan was 20 (Figure 2). As expected, overall *π* was less reactive to the bottleneck. 50 ticks after the bottleneck decreased population size by 10, *π* had decreased by 0.9% when average lifespan was 1 (annual simulation), 0.3% when average lifespan was 2, 0.1% when average lifespan was 5, 0.07% when average lifespan was 10, and 0.04% when average lifespan was 20 (Figure 2). As expected, we also saw that the most severe bottlenecks resulted in the fastest loss of genetic diversity. In the simulations where average lifespan was 20, *θ*_*W*_ had decreased by 16.1% when the bottleneck intensity was 100, 2.5% when the bottleneck intensity was 10, and 0.3% when the bottleneck intensity was 2 by 50 ticks after the bottleneck.

**Figure 2.**
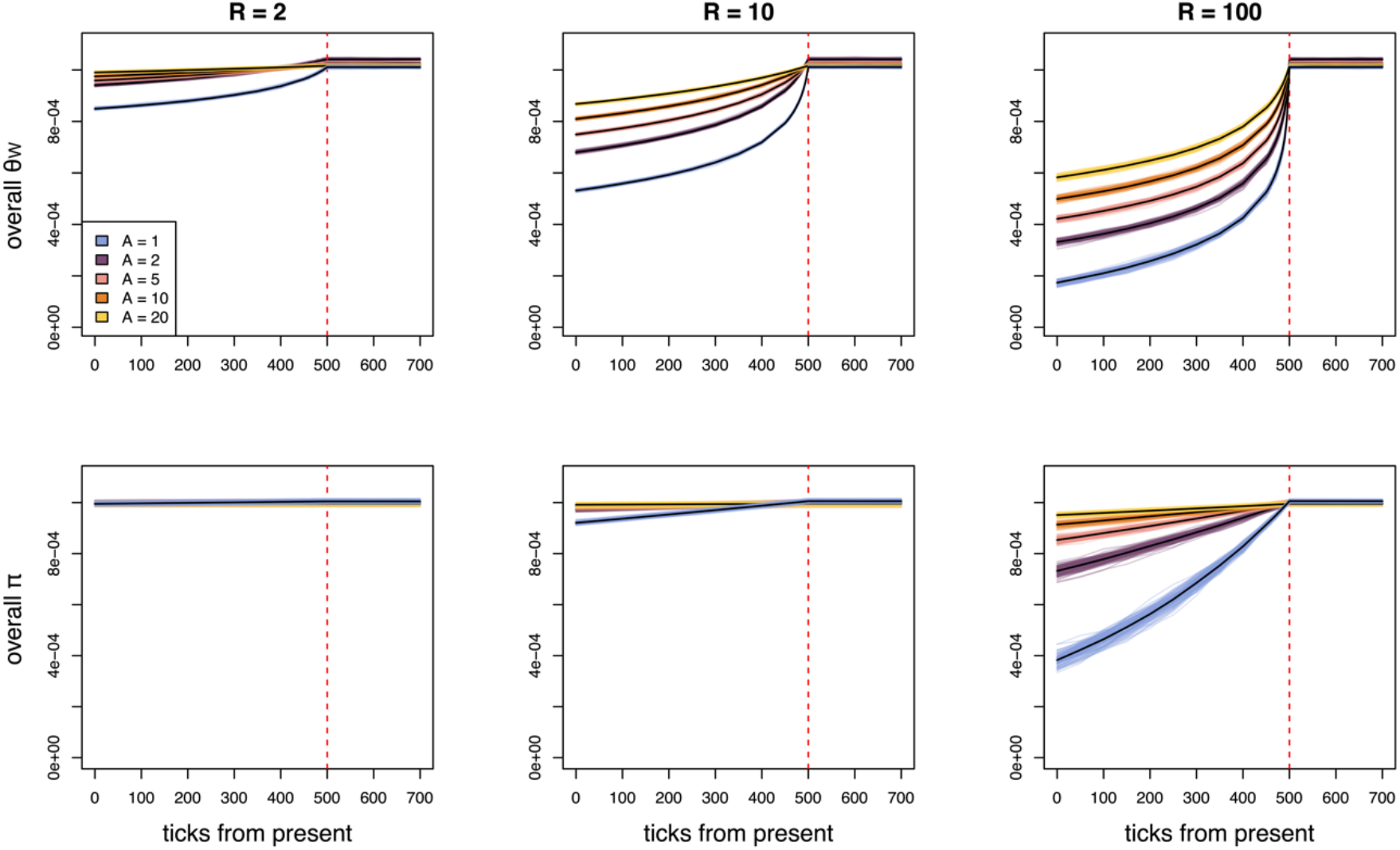
Plots show *θ*_*W*_ and *π* from all sampled individuals across time (ticks) for the perennial (average age, *A* = 2, 5, 10, 20) and annual (*A* = 1) simulations with three bottleneck severities (*R*, bottleneck severity increases with *R*). Colored lines represent different simulation replicates and black lines are the mean values across replicates. The red dashed line indicates when the bottleneck occurred.

**Figure 3.**
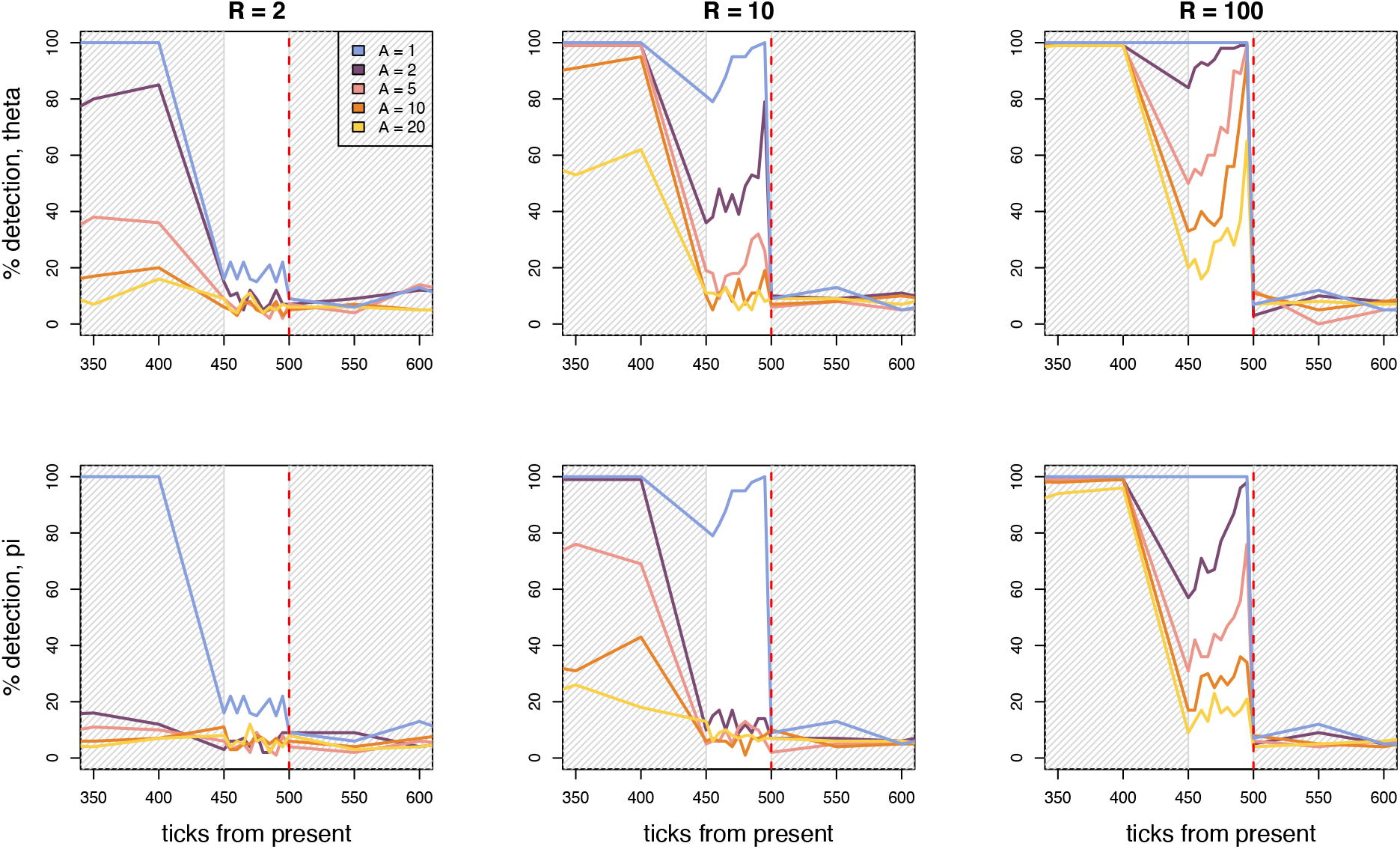
Plots show percent detection using temporal sampling for annual model (*A* = 1) and perennial models (*A* = 2, 5, 10, 20) for three bottleneck severities (*R*, bottleneck severity increases with *R*). Percent detection is defined as the percent of replicates where we found a significant difference between individuals subsampled from the present and previous timepoints. The red dashed line indicates when the bottleneck event occurred. The gray shaded portions of the plot indicate when samples were taken every 50 ticks of the simulation, whereas the unshaded portion of the plot indicates when samples were taking every five ticks of the simulation.

### 3.2. Temporal sampling

When comparing diversity from before and immediately after the R = 10 bottleneck event, we detected a change in *θ*_*W*_ in 100% of replicates when average lifespan was 1, 79% of replicates when average lifespan was 2, 26% of replicates when average lifespan was 5, 19% of replicates when average lifespan was 10, and 8% of replicates when average lifespan was 20 (Figure 3). Percent detection of bottlenecks increased with increasing bottleneck intensity. As expected, directly after the bottleneck, temporal comparison separated by five ticks have more power to detect the bottleneck in the annual model compared to the perennial models (Figure 3). Under severe bottleneck scenarios, temporal comparison performs similarly in annual and perennial models when the timepoints being compared were 50 ticks apart.

### 3.3. The utility of age information

To assess the relationship between genetic patterns and age information after a bottleneck, we evaluated the correlation between age and *θ*_*W*_, and age and *π* under a scenario where individual age can only be approximated. We compared *θ*_*W*_ and *π* between the oldest and youngest individuals at each sampled timepoint and assessed significance compared to a randomly permuted distribution of differences (Figure 4). The strength of the bottleneck strongly influenced the differences in *θ*_*W*_ and *π* between the binned age groups. When average lifespan was 20, we found a significant difference between old and young individuals in 96% (*R* = 100), 13% (*R* = 10), and 4% (*R* = 2) of replicates five ticks after the bottleneck. Fifty ticks after the bottleneck, we found significant differences in 42% (*R* = 100), 15% (*R* = 10), and 13% (*R* = 2) of replicates for the same simulations (Figure 4). When we expand the older age bin to include individuals above the 50^th^ quantile of ages, detection is similar across bottleneck severities five ticks after the bottleneck (*R* = 100: 95%, *R* = 10: 13%, *R* = 2: 4%), but decreases by 50 ticks after the bottleneck (*R* = 100: 53%, *R* = 10: 7%, *R* = 2: 9%, Supplemental Figure 1).

**Figure 4.**
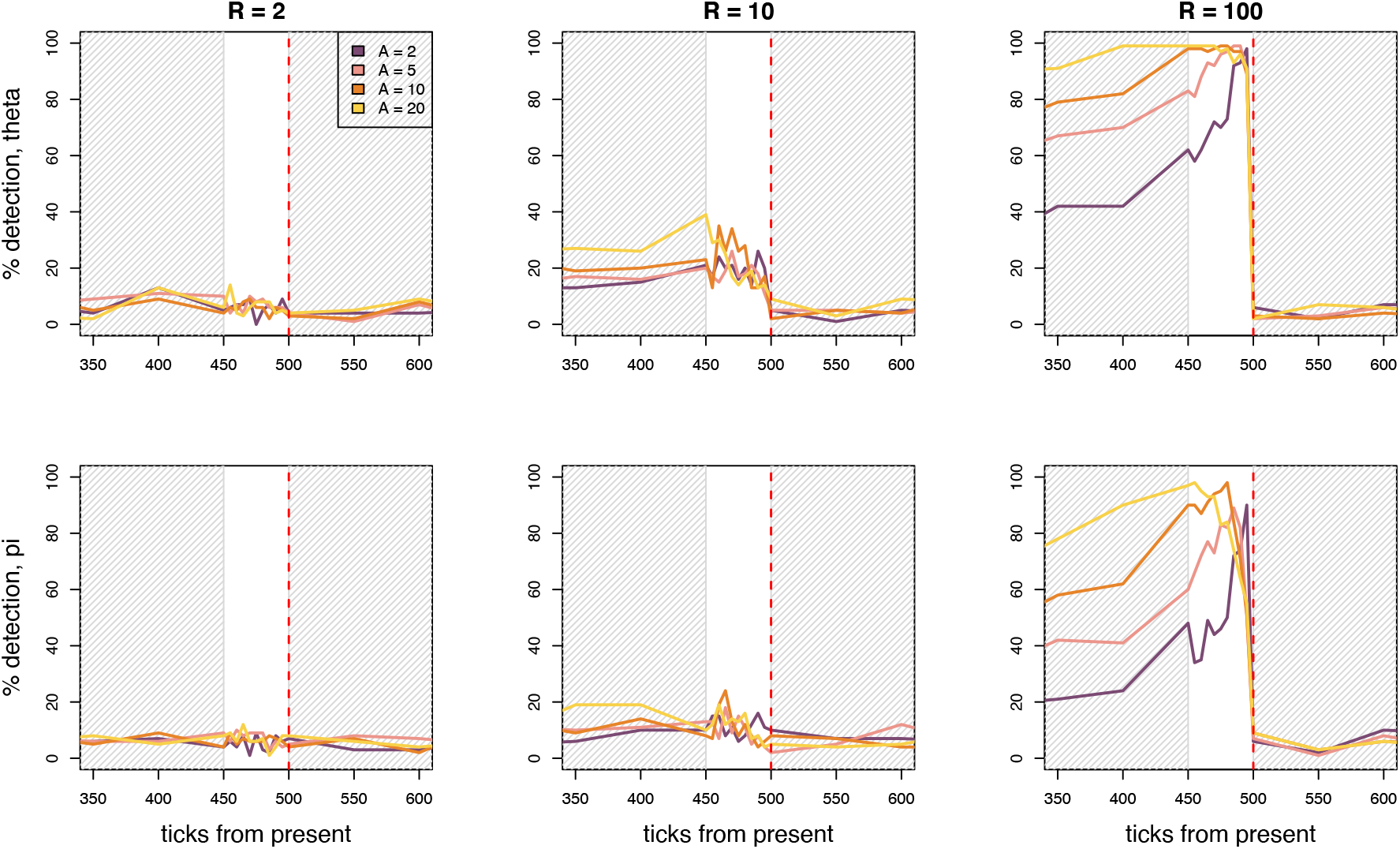
Plots show percent detection in *θ*_*W*_ and *π* between old and young individuals for perennial models (*A* = 2, 5, 10, 20). The old age bin is made up of individuals whose age at a given timepoint fell above the 90^th^ quantile. The young age bin is made up of the youngest individuals at a given timepoint. Sample size is the same between bins. Percent detection is defined as the percent of replicates where we found a significant difference between the old and young age bins. The red dashed line indicates when the bottleneck event occurred. The gray shaded portions of the plot indicate when samples were taken every 50 ticks of the simulation, where the unshaded portion of the plot indicates when samples were taking every five ticks of the simulation.

To compare the relative power of temporal sampling and age bin sampling, we calculated the difference in detection between temporal and age bin sampling methods (Figure 5). When the bottleneck was most severe (*R* = 100), age bin sampling out-performed temporal sampling for the *A* = 5, 10 and 20 simulations, but not when *A* = 2. Temporal sampling and age bin sampling performed similarly at lower bottleneck intensities (*R* = 2, *R* = 10).

**Figure 5.**
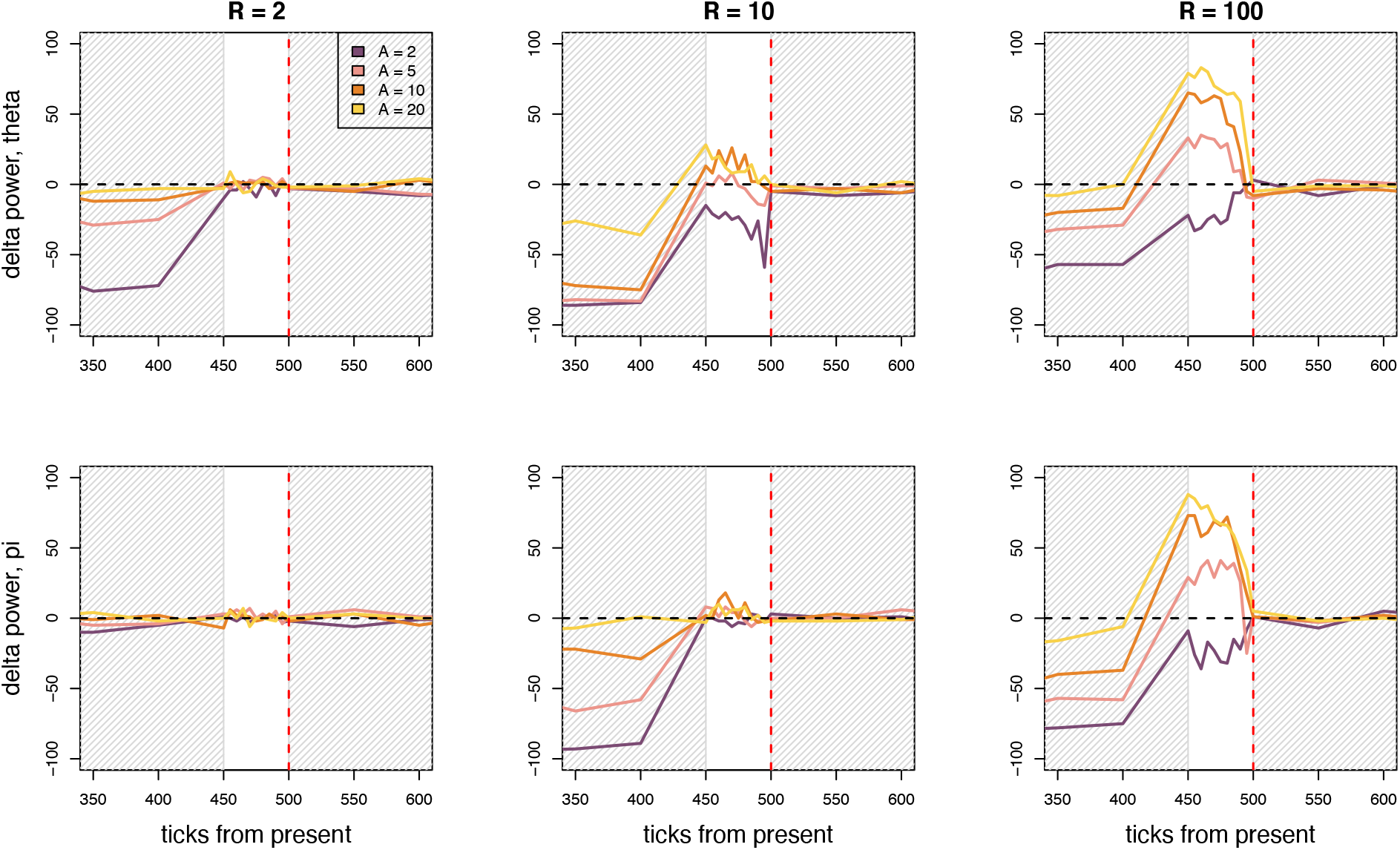
Plots show the difference in detection power between temporal and age bin sampling methods. Positive values indicate that age bin methods had higher percent detection and negative values indicate that temporal methods had higher percent detection. The red dashed line indicates when the bottleneck event occurred. The gray shaded portions of the plot indicate when samples were taken every 50 ticks of the simulation, where the unshaded portion of the plot indicates when samples were taking every 5 ticks of the

## 4. Discussion

### 4.1. Use of age information in detecting declines

Solving the biodiversity crisis necessitates the clever use and exploration of data from imperiled species, and our results highlight that age information could be a valuable tool when assessing the recent demography of long-lived species. In this study, we demonstrate the impact of generational overlap and longevity on genetic diversity and test the utility of age information when detecting demographic declines in species with overlapping generations. In accordance with theory, we found that the rate at which diversity was lost after a bottleneck event decreased with increasing average lifespan in our simulations (Figure 2). Also as expected, we find that *θ*_*W*_ is more sensitive to declines than *π* (Figure 2). We simulated temporal sampling by comparing diversity between adjacent timepoints and found that percent detection of the bottleneck event decreased with increasing lifespan. Temporal sampling before and after a bottleneck event is unlikely to be available for most populations. Our results suggest that age sampling––comparing younger and older individuals within a population, could support the inference of a recent bottleneck from a single sampling point, but that its use is limited when a decline has been less severe (*R* = 2, 10). In general, age comparisons outperformed temporal tests when the generation time in a simulation was ≥ five and when the bottleneck was severe.

### 4.2. Considerations for empirical applications

Age structure is being recognized as an important source of information about long-lived species (Dolan et al., 2023; Holmes & York, 2003). Although age information is rarely readily available for empirical populations, it is sometimes possible to approximate an individual’s age from their phenotype (e.g., turtle scutes and carapace length (Jensen et al., 2018; Wilson et al., 2003), rattlesnake rattles (Heyrend & Call, 1951), tree rings (Shroder, 1980), otoliths (Campana, 1999), tooth wear (Hinton et al., 2023), telomere length (Haussmann & Vleck, 2002), DNA methylation (Nakamura et al., 2023), coloration (Pyle, 1997) etc.) or population monitoring (Eaton & Link, 2011)). Noninvasive aging methods seem particularly promising for populations of conservation concern.

Here, we find that even having approximate age information increases power for detecting patterns of severe decline compared to temporal sampling, though increasing the uncertainty around adult age classes by expanding the older age bin decreases detection (Supplemental Figure 1). Using individuals binned by age ranges (as opposed to exact age information) might be superior to precise age information, as small sample sizes within each age cohort will limit inference and binning individuals based on approximate age allows for comparison between larger groups. For example, *θ*_*W*_ is biased at small sample sizes under non-Wright-Fisher conditions, so care is needed when interpreting *θ*_*W*_ calculated from only a few individuals.

Based on our simulations, the ideal scenario to incorporate age information into analyses would be in populations of species with a generation time greater than 11 years, where there is reason to believe a significant demographic decline has occurred, where methods are available to approximate age, ideally including estimates of age into adulthood, and where adequate sample sizes are available. Given our low detection probabilities using age comparisons under less severe bottleneck intensities (*R = 2, 10*), our results also highlight that a lack of a relationship between age and diversity in empirical data is not necessarily evidence that no decline has occurred, which could explain studies that have not found a relationship between age and genetic diversity in putatively declining populations (Schmidt et al., 2018).

Simulating data allowed us to test questions about age and genetic patterns in a controlled fashion. We had access to error-free whole-chromosome sequences for our samples and were only constrained by computational limitations. In wild populations, DNA quality, type of sequencing, and data quality will impact the power of inference. Similarly, we simulated populations isolated from the effects of gene flow. Immigrants from populations with different demographic trajectories are likely to complicate comparisons between older and younger individuals. Additionally, we simulated a bottleneck event that occurred instantaneously. Empirical examples of such declines include natural disasters and disease outbreaks (Hsu et al., 2017; McCallum, 2012), however many actual declines are gradual and occur on the timescale of decades (although may also be of much higher magnitude). Future research should investigate the use of age sampling when declines occur over a longer period.

Selection dynamics too will alter patterns seen in empirical systems. Inbreeding depression can result in viability selection, where adults in a population have higher heterozygosity because low-heterozygosity individuals do not reach adulthood due to deleterious homozygous genotypes (Clegg & Allard, 1973; Doyle et al., 2019; Labonne et al., 2016). Here we show that adults can have higher genetic diversity compared to juveniles due to purely neutral processes if there has been a recent population decline. It is possible that viability selection and neutral processes after a bottleneck could have an additive effect on the diversity differences between age groups.

Many different calculations of N_e_ exist; these are reflective of different attributes of the focal population, or of the focal population at varying points in the past (Nadachowska-Brzyska et al., 2022). In this study, we use *θ*_*W*_ and *π* as reflections of long-term N_e_. Calculations of N_e_ more sensitive to contemporary changes in population size, like patterns of linkage disequilibrium and allele frequency changes over time, should hold even more power to detect recent declines (England et al., 2010; Hollenbeck et al., 2016; Nadachowska-Brzyska et al., 2022; Waples, 2022; Waples & Do, 2008). Although computationally infeasible within our experimental design, if appropriate genomic data is available for the population of concern, it may be fruitful to incorporate linkage information into comparisons between age bins. We also note that we may have had higher power to detect declines with more advanced demographic inference methods that can incorporate overlapping generations (Kamm et al., 2020), but to our knowledge, there are no demographic inference methods that incorporate the age of sampled individuals, though this could be an intriguing avenue of future development.

### 4.3. Life history impacts patterns of neutral genetic diversity

Our results highlight that the life history of an organism is intrinsically linked to N_e_ and to neutral genetic patterns at the individual and population level. Common model assumptions, like that of Wright-Fisher models, are violated by long-lived species in ways that impact our interpretation of data, particularly when inferring changes in N_e_. Because evolution in long-lived species moves at a slower pace compared to shorter lived species (Bergeron et al., 2023), they may be more at risk of being outpaced by shifting climatic conditions and landscape changes. However, these species also maintain higher N_e_ compared to species with shorter generation times after declines, perhaps allowing for conservation intervention to rescue populations before they enter an extinction vortex (Gilpin & Soulé, 1986). In the absence of temporal sampling, our simulations suggest that age information is an additional tool researchers can use in detecting recent demographic bottlenecks in long-lived species.

## Supporting information

Supplemental materials: perennial model

Supplemental materials: annual model

## Acknowledgements

All simulations and analyses were run on the High Performance Computing Center at Michigan State University’s Institute for Cyber-enabled Research. We would like to thank Bob Week, Matteo Tomasini, and Zach Hancock for help and feedback on the development of the simulations, Peter Ralph for guidance using pyslim, Tyler Linderoth for fruitful discussion on the true nature of theta, and members of the Bradburd, Fitzpatrick, and Winger Labs for valuable feedback on this paper. Research reported in this publication was supported by the National Institute of General Medical Sciences of the National Institutes of Health under Award Number R35GM137919 (awarded to GB). The content is solely the responsibility of the authors and does not necessarily represent the official views of the NIH. This is W. K. Kellogg Biological Station Contribution No. 2374.

## Data Accessibility

Code for all simulations and scripts for all downstream analyses will be available on a dryad repository (https://datadryad.org/).

